# Characterization of SARS-CoV-2 Omicron BA.2.75 clinical isolates

**DOI:** 10.1101/2022.08.26.505450

**Authors:** Ryuta Uraki, Shun Iida, Peter J. Halfmann, Seiya Yamayoshi, Yuichiro Hirata, Kiyoko Iwatsuki-Horimoto, Maki Kiso, Mutsumi Ito, Yuri Furusawa, Hiroshi Ueki, Yuko Sakai-Tagawa, Makoto Kuroda, Tadashi Maemura, Taksoo Kim, Sohtaro Mine, Noriko Iwamoto, Rong Li, Yanan Liu, Deanna Larson, Shuetsu Fukushi, Shinji Watanabe, Ken Maeda, Zhongde Wang, Norio Ohmagari, James Theiler, Will Fischer, Bette Korber, Masaki Imai, Tadaki Suzuki, Yoshihiro Kawaoka

**Affiliations:** Division of Virology, Institute of Medical Science, University of Tokyo, Tokyo 108-8639, Japan; The Research Center for Global Viral Diseases, National Center for Global Health and Medicine Research Institute, Tokyo 162-8655, Japan; Department of Pathology, National Institute of Infectious Diseases, Tokyo 162-8640, Japan; Influenza Research Institute, Department of Pathobiological Sciences, School of Veterinary Medicine, University of Wisconsin-Madison, Madison, WI 53711, USA; Disease Control and Prevention Center, National Center for Global Health and Medicine Hospital, Tokyo 162-8655, Japan; Department of Animal, Dairy, and Veterinary Sciences, College of Agriculture and Applied Sciences, Utah State University, Logan, UT 84322, USA; Department of Virology 1, National Institute of Infectious Diseases, Musashimurayama, Tokyo 208-0011, Japan; Center for Influenza and Respiratory Virus Research, National Institute of Infectious Diseases, Musashimurayama, Tokyo 208-0011, Japan; Department of Veterinary Science, National Institute of Infectious Diseases, Tokyo 162-8640, Japan; Space Data Science and Systems, Los Alamos National Laboratory, Los Alamos, NM 87545, USA; Theoretical Biology and Biophysics, Los Alamos National Laboratory, Los Alamos, NM 87545, USA; New Mexico Consortium, Los Alamos, NM 87545, USA

**Keywords:** BA.2.75, Omicron, Syrian hamster, hACE2-expressing hamster, lung inflammation

## Abstract

The prevalence of the Omicron subvariant BA.2.75 is rapidly increasing in India and Nepal. In addition, BA.2.75 has been detected in at least 34 other countries and is spreading globally. However, the virological features of BA.2.75 are largely unknown. Here, we evaluated the replicative ability and pathogenicity of BA.2.75 clinical isolates in Syrian hamsters. Although we found no substantial differences in weight change among hamsters infected with BA.2, BA.5, or BA.2.75, the replicative ability of BA.2.75 in the lungs was higher than that of BA.2 and BA.5. Of note, BA.2.75 caused focal viral pneumonia in hamsters, characterized by patchy inflammation interspersed in alveolar regions, which was not observed in BA.5-infected hamsters. Moreover, in competition assays, BA.2.75 replicated better than BA.5 in the lungs of hamsters. These results suggest that BA.2.75 can cause more severe respiratory disease than BA.5 and BA.2 and should be closely monitored.

## Introduction

Severe acute respiratory syndrome coronavirus 2 (SARS-CoV-2), first detected in China at the end of 2019, is responsible for COVID-19, which is associated with mild to severe symptoms ranging from cough and fever to severe pneumonia and death. Over two years have passed since the World Health Organization (WHO) declared COVID-19 a pandemic (https://covid19.who.int/). Yet, SARS-CoV-2 still imposes huge public health and economic burdens worldwide. The currently circulating Omicron (B.1.1.529) variant emerged at the end of 2021 and has since evolved into complex sublineages; three of the major Omicron lineages have serially transitioned as globally dominant forms: first BA.1, then BA.2, and then BA.5 (**Fig. 1a**). The BA.5 lineage is currently the dominant variant circulating globally (https://covariants.org/per-variant). BA.5 was just beginning to expand in India in May 2022, when BA.2.75 (a subvariant of the BA.2 sublineage) first emerged there. This subvariant appears to be more transmissible than BA.5 in India and Nepal, where it is gaining prevalence (https://covariants.org/per-variant). Recently, WHO has categorized BA.2.75 as a variant of concern (VOC) lineages under monitoring (VOC-LUM).

**Fig. 1.**
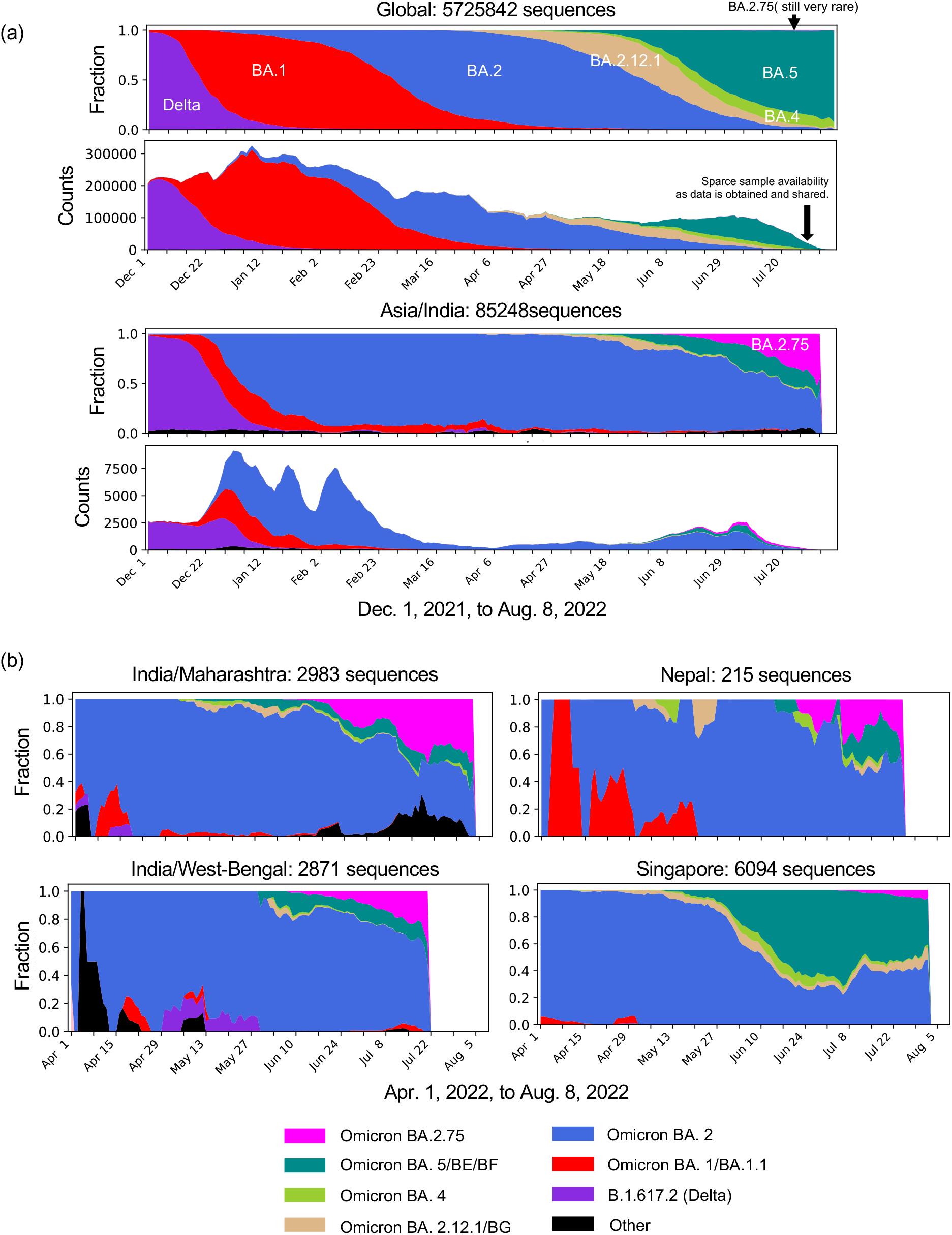
Pango lineage dynamics between 01-12-2021 and 08-08-2022. **a**, BA.2.75 frequencies over time globally and in India, where BA.2.75 is currently most commonly found. Omicron variants have been through waves of global dominance since Omicron began to spread in late 2021. BA.1 (and BA.1.1, both in red) very rapidly replaced Delta globally. BA.2 then replaced BA.1 as the globally dominant form. Within the BA.2 lineage, BA.2.12.1 began to expand in North America, but BA.2, including the BA.2.12.1 sublineage, is being replaced by BA.5, the currently globally dominant form. BA.2.75 is still very rare globally, at only 0.3% of the global sample in the 30 days ending on 08-08-2022, but is being increasingly sampled in India, representing 25% of the last 30-day sample. BA.5 was slower to begin its expansion in India than in most countries, but was still on a trajectory of increasing prevalence when BA.2.75 was first sampled in late May/early June; BA.2.75 is much more rapidly gaining in prevalence in India than BA.5, suggesting a possible selective advantage. **b**, Examples illustrating the increase in prevalence of BA.2.75 relative to BA.5, despite BA.5 being well-established prior to BA.2.75’s introduction, at both the state level within India (left), and in countries outside of India (right). Singapore and Nepal were selected as examples from Fig. S2 as they were the two countries with the highest frequency of BA.2.75 outside of India. Maharashtra and West Bengal were selected from Sup. Fig. S3 as they are the states within India that currently have the most available samples. All data are from GISAID; the illustrations we made with the “Embers” web-based tool at cov.lanl.gov (Korber et al., 2020).

Compared with the original Wuhan Hu-1 strain, the Omicron BA.1 virus had more than 30 amino acid differences in the spike protein of SARS-CoV-2 including insertions and deletions (**Fig. S1A**) (by comparison, Delta differed from the original Wuhan Hu-1 by only 11 amino acids in its Spike)(Flemming, 2022). BA.2 differed from BA.1 at 27 Spike positions, and BA.5 differs from BA.2 by 5 amino acids in the S protein (**Fig. S1A**). We recently demonstrated that the pathogenicity of BA.1 and BA.2 sublineage viruses is comparable in animal models and attenuated compared with previously circulating variants of concern (VOCs), consistent with clinical data in humans (Halfmann et al., 2022; Uraki et al., 2022a). In addition, our recent data suggest that BA.4 and BA.5 have similar pathogenicity to that of BA.2 in rodent models ((Kawaoka et al., 2022): https://www.researchsquare.com/article/rs-1820048/v1). SARS-CoV-2 initiates infection through the binding of the receptor-binding domain (RBD) of its spike protein to host cell surface receptors [i.e., human angiotensin-converting enzyme 2 (hACE2)]. BA.2.75 differs from that of BA.2 by nine amino acids in the Spike, including four in the RBD (i.e., G339H, G446S, N460K, and the wild-type amino acid at position Q493). Recent studies reported that the RBD of BA.2.75 has a higher binding affinity for hACE2 than that of BA.2 ((Cao et al., 2022): https://www.biorxiv.org/content/10.1101/2022.07.18.500332v1.full, (Saito et al., 2022): https://www.biorxiv.org/content/10.1101/2022.08.07.503115v1), raising the possibility that this property may increase the replicative ability and/or pathogenicity of BA.2.75. Moreover, in addition to the substitutions in the RBD, there are several amino acid differences in the other viral proteins of BA.2.75, which may also alter its replicative capability and pathogenicity (**Fig. S1b**). Here, we assessed the replicative capacity and pathogenicity of authentic BA.2.75 subvariants isolated from COVID-19 patients in established COVID-19 animal models.

## Results

### Transitions in Omicron variant prevalence throughout 2022

SARS-CoV-2 has undergone a series of variant transitions since the Omicron lineage was first observed in November of 2021. The initial global transition from Delta to the Omicron BA.1 lineage was extremely swift and was followed successively by waves of BA.2 and BA.5, with each variant, essentially replacing the previous dominant form (**Fig. 1**). This may indicate that each variant has been more transmissible than the prior variant, particularly in settings with histories of prior infection and vaccination resulting in changes in immune status at the population level. BA.2.75, a BA.2 sublineage, was first detected in India in May of 2022, and since then has been rapid increasing in sampling frequency (**Fig.1, Fig. S2 and S3**). As of this writing, there are more than 3,000 sequences in GISAID (Elbe and Buckland-Merrett, 2017; Khare et al., 2021) with the Pango lineage designation BA.2.75, sampled in 35 nations. Although BA.2.75 is still rare outside of India and Nepal, it has been sampled at least 10 times in 10 nations, and in each of these 10 it is significantly increasing in sampling frequency (**Fig. S2**); the regularity of this pattern suggests that BA.2.75 may have a selective advantage over co-circulating variants. BA.2.75 is established in 12 states in India, and is significantly increasing in frequency throughout India, indicating that its increased prevalence in India is unlikely to be a founder effect or sampling issue (**Fig. S3**). BA.5 was just beginning to expand in India and when BA.2.75 was first detected (**Fig. 1**), and where BA.5 and BA.2.75 are co-circulating, the prevalence of BA.2.75 tends to be increasing faster (**Fig. 1b**). Therefore, BA.2.75 is a likely candidate for the next major transition to a more transmissible form, unless a novel variant emerges with an even greater selective advantage.

### BA.2.75 infection in hamsters

To characterize BA.2.75 *in vivo*, we amplified three BA.2.75 clinical isolates in VeroE6/TMPRSS2 cells: hCoV-19/Japan/TY41-716/2022 (TY41-716)(Takashita et al., 2022), hCoV-19/Japan/UT-NCD1757-1N/2022 (NCD1757), and hCoV-19/Japan/UT-NCD1759-1N/2022 (NCD1759). We confirmed that the S protein of all three isolates contained the nine additional amino acid changes (i.e., K147E, W152R, F157L, I210V, G257S, D339H, G446S, N460K, and Q493 (reversion)) (**Fig. S1b**) that distinguish the consensus form of BA.2.75 (https://cov.lanl.gov/components/sequence/COV/pangocommonforms.comp) from a BA.2 isolate (hCoV-19/Japan/UT-NCD1288-2N/2022; NCD1288), which carries the most common circulating form of BA.2 in Spike. However, two of the isolates (NCD1757 and NCD1759) had a D574V substitution in the subdomain (SD), in addition to the nine mutations; this and several other distinctive mutations found in other proteins are summarized in **Fig. S1b**.

We first evaluated the pathogenicity of the BA.2.75 isolates in wild-type Syrian hamsters, a well-established small animal model for the study of COVID-19 (Chan et al., 2020; Imai et al., 2020; Sia et al., 2020). Syrian hamsters were intranasally inoculated with 10^5^ plaque-forming units (PFU) of BA.2.75 (TY41-716, NCD1757, or NCD1759). For comparison, additional hamsters were infected with clinical isolates of BA.2 (10^5^ PFU of NCD1288)(Takashita et al., 2022; Uraki et al., 2022a), BA.5 [10^5^ PFU of hCoV-19/Japan/TY41-702/2022 (TY41-702)] (Kawaoka et al., 2022), or B.1.617.2 [10^5^ PFU of hCoV-19/USA/WI-UW-5250/2021 (Delta: UW5250)](Halfmann et al., 2022). Intranasal infection with B.1.617.2 resulted in significant body weight loss by 6 days post-infection (dpi) (-5.4%) (**Fig. 2a**), consistent with our previous observations (Halfmann et al., 2022; Kawaoka et al., 2022). By contrast, most of the animals infected with any of the three BA.2.75 isolates gained weight over the 6-day experiment, similar to BA.2-, BA.5-, or mock-infected animals. We also examined pulmonary functions in the infected hamsters by measuring Penh and Rpef, which are surrogate markers for bronchoconstriction and airway obstruction, respectively, by using a whole-body plethysmography system. Inoculation of hamsters with the BA.2, BA.5, BA.2.75 (NCD1757), or BA2.75 (NCD1759) isolate did not cause substantial changes in either Penh or Rpef at any timepoint post-infection compared to the mock-infected group. Infection with BA.2.75 (TY41-716) caused a slight increase in Penh at 3 and 5 dpi, although no statistically significant differences in Penh values were observed among BA.2-, BA.5-, and BA.2.75 (TY41-716)-infected animals. Consistent with our previous data, infection with B.1.617.2 caused significant changes in Rpef in comparison with the five Omicron isolates (**Fig. 2b**).

**Figure 2.**
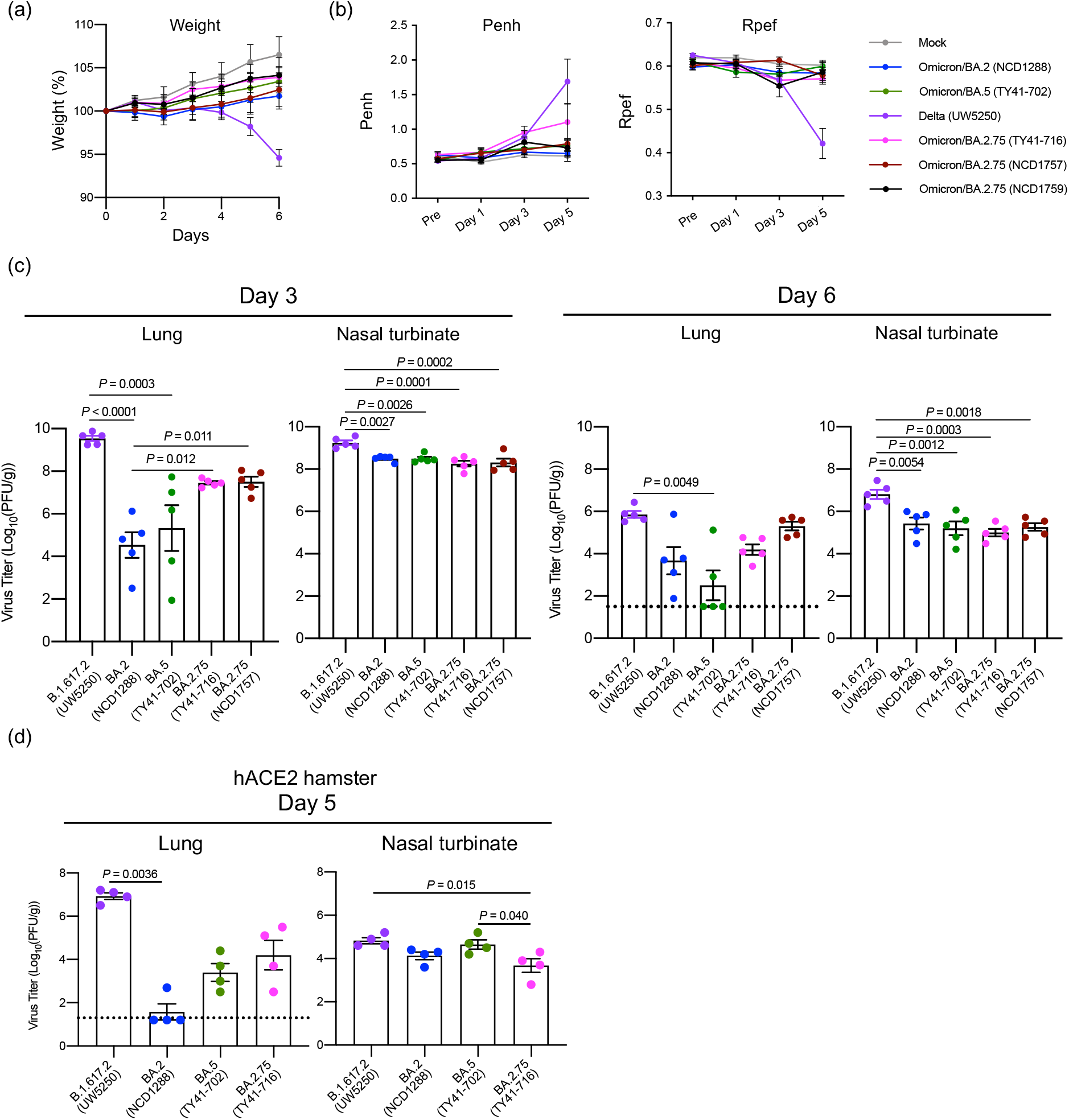
The infectivity and pathogenicity of BA.2.75 in hamsters. **a**,**b**, Wild-type Syrian hamsters were intranasally inoculated with 10^5^ PFU in 30 μL of BA.2 (NCD1288) (n=9), BA.5 (TY41-702) (n=9), BA.2.75 (TY41-716) (n=5), BA.2.75 (NCD1757) (n=5), BA.2.75 (NCD1759) (n=5), B.1.617.2 (UW5250) (n=9), or PBS (mock) (n=8). **a**, Body weights of virus-infected and mock-infected hamsters were monitored daily for 6 days. Data are presented as the mean percentages of the starting weight (± s.e.m.). **b**, Pulmonary function analyses in virus-infected and mock-infected hamsters. Penh and Rpef were measured by using whole-body plethysmography. Mean ± s.e.m. Data were analyzed by using a two-way ANOVA followed by Tukey’s multiple comparisons test. **c**, Virus replication in infected Syrian hamsters. Hamsters (*n* =10) were intranasally inoculated with 10^5^ PFU in 30 μL of BA.2 (NCD1288), BA.5 (TY41-702), BA.2.75 (TY41-716), BA.2.75 (NCD1757), or B.1.617.2 (UW5250) and euthanized at 3 and 6 dpi for virus titration (*n* =5/day). Virus titers in the nasal turbinates and lungs were determined by performing plaque assays with Vero E6-TMPRSS2-T2A-ACE2 cells. Vertical bars show the mean ± s.e.m. Points indicate data from individual hamsters. The lower limit of detection is indicated by the horizontal dashed line. Data were analyzed by using a one-way ANOVA with Tukey’s multiple comparisons test (titers in the lungs at 3 dpi and nasal turbinates at 3 and 6 dpi) or the Kruskal-Wallis test followed by Dunn’s test (titers in the lungs at 6 dpi). **d**, hACE2-expressing Syrian hamsters (*n* =4) were intranasally inoculated with 10^5^ PFU in 30 μL of BA.2 (NCD1288), BA.5 (TY41-702), BA.2.75 (TY41-716), or B.1.617.2 (UW5250). Infected animals were euthanized at 5 dpi for virus titration (*n* =4/group). Virus titers in the nasal turbinates and lungs were determined by performing plaque assays with Vero E6-TMPRSS2-T2A-ACE2 cells. Vertical bars show the mean ± s.e.m. Points indicate data from individual animals. The lower limit of detection is indicated by the horizontal dashed line. Data were analyzed by using a one-way ANOVA with Tukey’s multiple comparisons test (titers in the nasal turbinates) or the Kruskal-Wallis test followed by Dunn’s test (titers in the lungs). *P* values of < 0.05 were considered statistically significant.

We next assessed levels of infection in the respiratory tract of wild-type Syrian hamsters (**Fig. 2c**). Hamsters were intranasally infected with 10^5^ PFU of BA.2.75 (TY41-716), BA.2.75 (NCD1757), BA.2 (NCD1288), BA.5 (TY41-702), or B.1.617.2 (Delta: UW5250); at 3 and 6 dpi, the animals were sacrificed, and their nasal turbinates and lungs were collected for virus titration. The virus titers were determined by performing plaque assays on Vero E6-TMPRSS2-T2A-ACE2 cells. BA.2 (NCD1288), BA.5 (TY41-702), BA.2.75 (TY41-716), and BA.2.75 (NCD1757) replicated in the nasal turbinates of the infected animals with no significant differences in viral titers at both timepoints examined. However, the virus titers in the nasal turbinates were significantly lower in the respiratory tract of animals infected with the BA.2, BA.5, BA.2.75 (TY41-716), or BA.2.75 (NCD1757) isolates, compared to animals infected with B.1.617.2 [mean differences in viral titer = 0.75, 0.75, 0.98, or 0.94 and 1.4, 1.6, 1.8, or 1.5 log_10_ (PFU/g) at 3 and 6 dpi, respectively].

Consistent with our previous report (Kawaoka et al., 2022), the virus titers in the lungs of animals infected with BA.2 or BA.5 were lower than those in animals infected with B.1.617.2 [mean differences in viral titer = 5.0 or 4.2 and 2.2 or 3.4 log_10_ (PFU/g) at 3 and 6 dpi, respectively], although the difference was not statistically significant between the BA.2- and B.1.617.2-infected groups at 6 dpi. The lung titers in the BA.2.75 (TY41-716)-infected groups were also lower than those in the B.1.617.2-infected groups [mean difference in viral titer = 2.1 and 1.7 log_10_ (PFU/g) at 3 and 6 dpi, respectively], although these differences did not reach statistical significance. The viral titers in the lungs of another BA.2.75 strain (NCD1757)-infected groups were similarly lower than those in the B.1.617.2-infected group at 3 dpi [mean differences in viral titer = 2.0 log_10_ (PFU/g)]; however, animals infected with BA.2.75 or B.1.617.2 had similar titers in the lungs at 6 dpi. The lung titers in the BA.2.75 (TY41-716)- and BA.2.75 (NCD1757)-infected groups were higher than those in BA.5-infected groups [for BA.2.75 (TY41-716), mean differences in viral titer = 2.1 and 1.7 log_10_ (PFU/g), at 3 and 6 dpi, respectively; for BA.2.75 (NCD1757), mean differences in viral titer = 2.2 and 2.8 log_10_ (PFU/g), at 3 and 6 dpi, respectively]; however, the differences were not statistically significant among the three groups. At 3 dpi, the virus titers in the lungs were significantly higher in the respiratory tract of animals infected with BA.2.75 (TY41-716), compared to animals infected with BA.2 (NCD1288) [mean difference in viral titer = 2.9 log_10_ (PFU/g)]; however, at 6 dpi, similar titers were detected in the lungs of animals inoculated with BA.2.75 (TY41-716) or BA.2. The viral titers in the lungs of the BA.2.75 (NCD1757)-infected groups were also higher than those in the BA.2 (NCD1288)-infected groups [mean differences in viral titer = 3.0 and 1.6 log_10_ (PFU/g), at 3 and 6 dpi, respectively], although the difference was not statistically significant between the BA.2.75 (NCD1757)- and BA.2-infected groups at 6 dpi. Taken together, these results suggest that the replicative ability of BA.2.75 in the lungs of wild-type hamsters is higher than that of previous Omicron variants, including BA.2 and BA.5.

We then investigated the infectivity of BA.2.75 in respiratory organs by using a more susceptible model, specifically transgenic hamsters expressing hACE2 (**Fig. 2d**). At 5 dpi, the virus titers in the lungs and nasal turbinates of hACE2-expressing hamsters infected with BA.2.75 (TY41-716) were lower than those in animals infected with B.1.617 (UW5250) [mean differences in viral titer = 2.7 and 1.1 log_10_ (PFU/g), respectively], although the differences in the lungs were not statistically significant between the two groups. Similar titers were detected in the lungs of animals inoculated with BA.2.75 (TY41-716) or BA.5 (TY41-702); however, the virus titers in the nasal turbinates of the animals infected with BA.2.75 were slightly but significantly lower than in those infected with BA.5 [mean differences in viral titer = 0.98 log_10_ (PFU/g)]. The virus titers in the lungs were substantially higher in the respiratory tract of animals infected with BA.2.75 (TY41-716) compared with animals infected with BA.2 (NCD1288) [mean differences in viral titer = 2.6 log_10_ (PFU/g)], although animals infected with BA.2.75 or BA.2 exhibited similar viral titers in nasal turbinates. These results suggest that BA.2.75 may have a higher replicative ability than BA.2 in the lungs of hACE2 transgenic hamsters.

### Histopathological findings in the lungs of SARS-CoV-2 BA.2.75 virus-inoculated Syrian hamsters

The lungs of Syrian hamsters that were inoculated with BA.2.75, BA.5, or B.1.617.2 were also analyzed histopathologically. Hamsters were intranasally inoculated with BA.2.75 (TY41-716), BA.5 (TY41-702), or B.1.617.2 (Delta, UW5250) and euthanized at 3 and 6 dpi for histopathological evaluation; representative images are shown in Figure 3.

**Figure 3.**
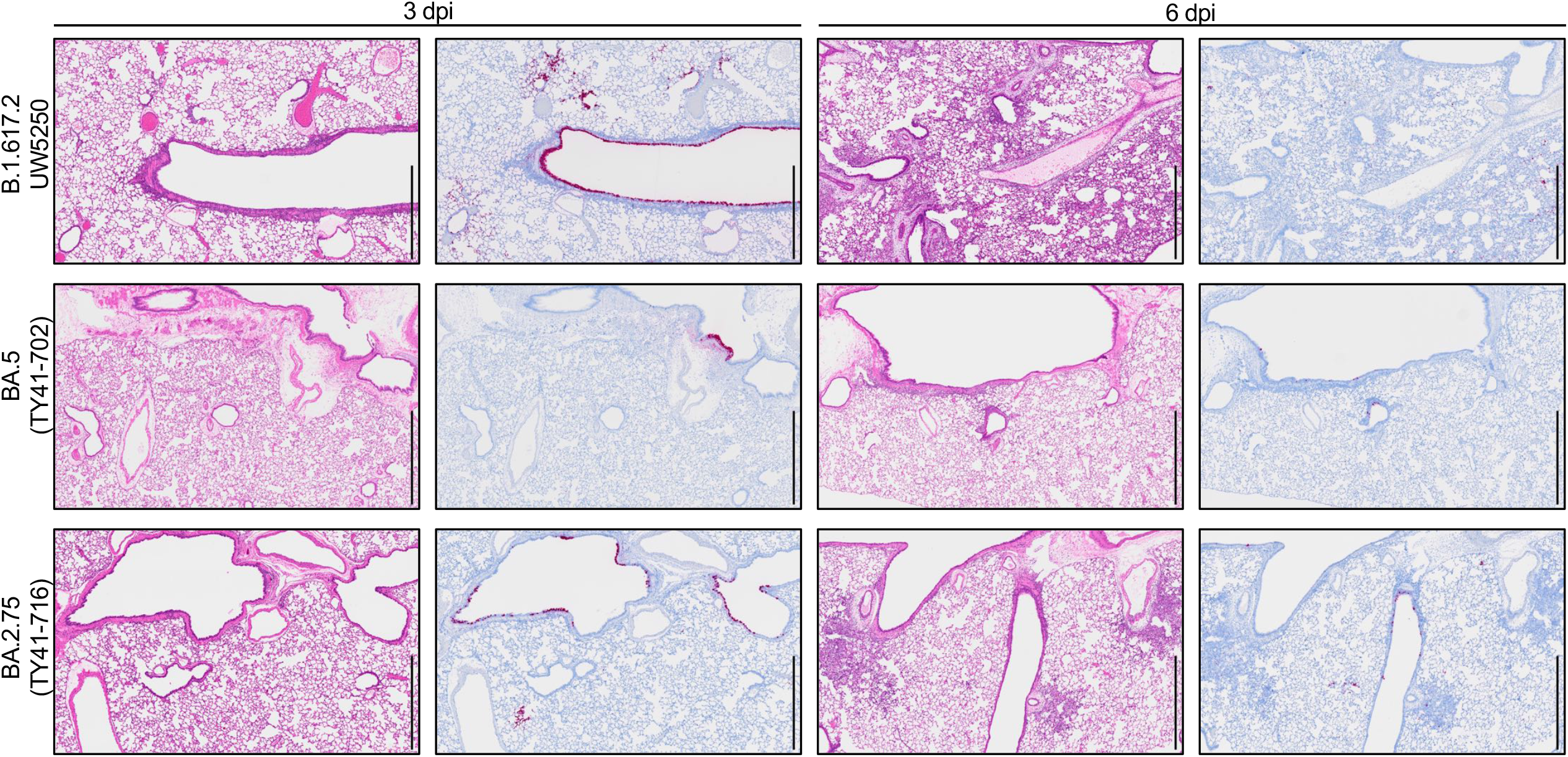
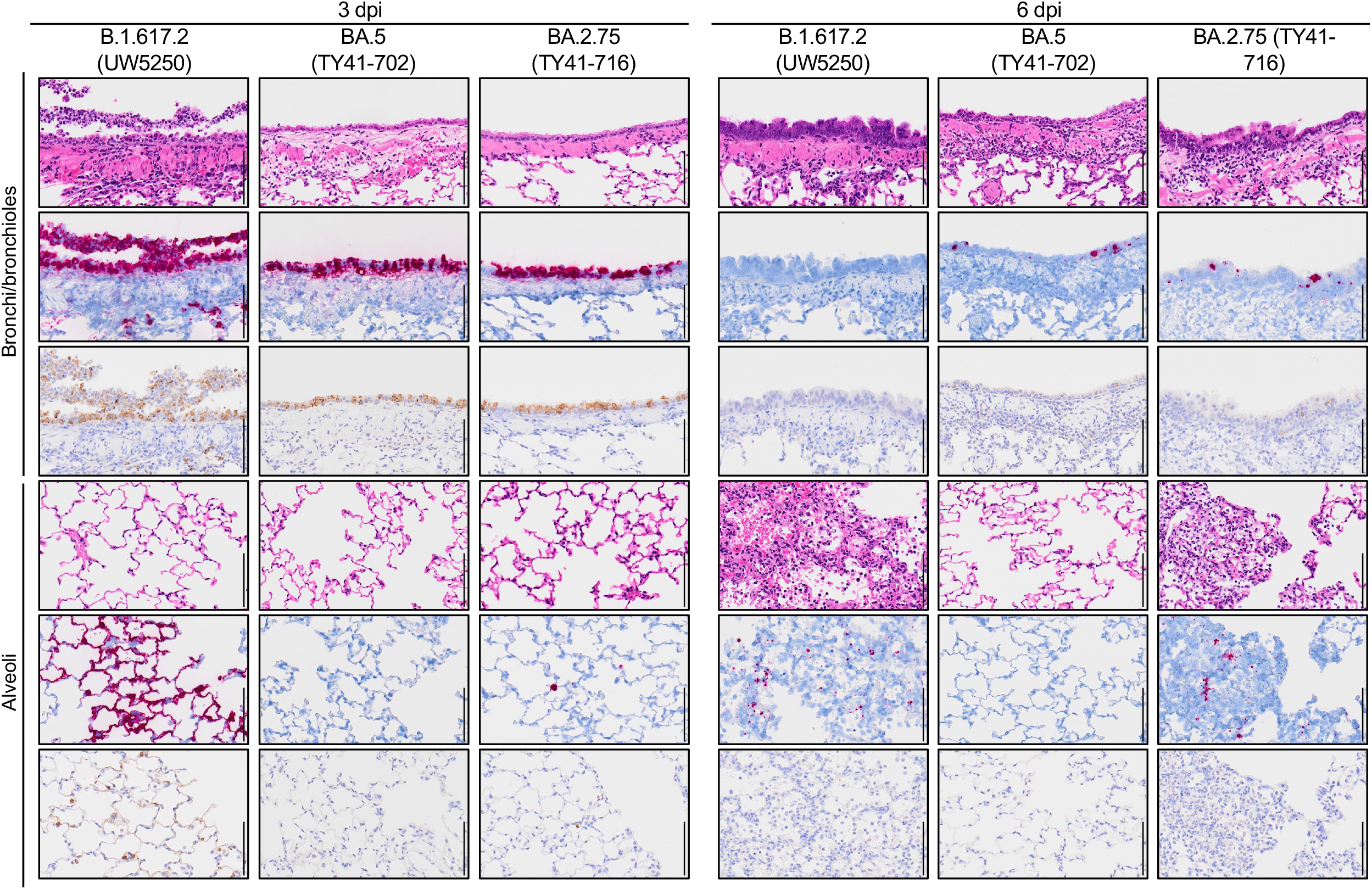
Histopathological findings in hamsters inoculated with BA.2.75. Syrian hamsters (n = 5, per group) were inoculated with 10^5^ PFU of BA.2.75 (TY41-716), BA.5 (TY41-702), or B.1.617.2 (UW5250) and sacrificed at 3 or 6 dpi for histopathological examinations. **a**, Representative images of the lungs at low magnification are shown. Left columns, hematoxylin and eosin staining. Right columns, in situ hybridization targeting the nucleocapsid gene of SARS-CoV-2. Scale bars, 1 mm. **b**, Representative images of the bronchi/bronchioles and alveoli at high magnification are shown. Upper rows, hematoxylin and eosin staining. Middle rows, in situ hybridization targeting the nucleocapsid gene of SARS-CoV-2. Lower rows: immunohistochemistry for the detection of SARS-CoV-2 nucleocapsid protein by a rabbit polyclonal antibody. Scale bars, 100 µm.

This examination revealed that inflammation was not obvious in the lungs of either BA.2.75 (TY41-716)- or BA.5-inoculated animals at 3 dpi; however, infiltration of inflammatory cells such as mononuclear cells and neutrophils was observed in peribronchial and peribronchiolar regions in these two groups at 6 dpi (**Fig. 3a, 3b, and S4**). It is noteworthy that focal pneumonia, characterized by patchy inflammation interspersed in alveolar regions, was observed in the lungs of BA.2.75 (TY41-716)-inoculated animals at 6 dpi. Similar histopathological findings (i.e., focal pneumonia) were observed in the lungs of animals inoculated with another BA.2.75 strain (NCD1757) at 6 dpi (**Fig. S5**). However, there was no obvious pneumonia in the lungs of BA.5-inoculated animals at the same timepoint. By contrast, in the lungs of the B.1.617.2-inoculated animals, peribronchial and peribronchiolar inflammation was prominent at 3 dpi, and extensive pneumonia with focal alveolar hemorrhage was observed in the alveolar regions at 6 dpi (**Fig. 3a, 3b and S4**). In addition, we detected viral RNA and protein in the lung tissue of BA.2.75 (TY41-716)-, BA.5- or B.1.617.2-infected hamsters by use of in situ hybridization and immunohistochemistry. These analyses revealed that viral RNA and antigen were readily detected on bronchial/bronchiolar epithelium in both BA.2.75 (TY41-716)- and BA.5-inoculated animals at 3 dpi with a clear decrease in positive cells over time (**Fig. 3a and 3b**). In the alveolar regions, a small number of cells were positive for viral RNA or antigen in the BA.2.75 (TY41-716)-inoculated group at both timepoints examined, and fewer cells were positive in the BA.5-inoculated group at the corresponding timepoints (**Fig. 3a and 3b**). Comparatively, at 3 dpi, the lungs of the B.1.617.2-inoculated hamsters had diffusely positive viral RNA and antigen in the bronchial/bronchiolar areas and patchily positive viral RNA and antigen in the alveolar regions (**Fig. 3a and 3b**). BA.2.75 (TY41-716) thus produced mild viral pneumonia in the hamster model with attenuated pathogenicity compared with B.1.617.2, whereas BA.5 did not cause obvious viral pneumonia. In addition, the number of viral RNA/antigen-positive cells in the alveolar regions of the BA.2.75 (TY41-716)-inoculated animals was higher than that in the BA.5-inoculated animals, but lower than that in the B.1.617.2-inoculated ones.

### The replicative fitness of BA.2.75 compared with that of BA.5 in hamsters

To further investigate the replicative fitness of BA.2.75, we compared the growth of BA.2.75 in wild-type hamsters with that of BA.5, which is currently the dominant variant circulating globally. Wild-type hamsters were intranasally inoculated with 2 × 10^5^ PFU of a mixture of BA.2.75 (TY41-716) and BA.5 (TY41-702) at ratios of 1:1, 1:3, 1:19, or 1:199. At 4 dpi, the proportion of each virus in the nasal turbinates and lungs of the infected hamsters was determined by using Next Generation Sequencing (NGS). The proportion was calculated on the basis of the differences between these two viruses across 6 regions in the S protein.

NGS analysis revealed that the proportion of BA.2.75 had increased in the nasal turbinates and lungs of all infected animals compared to that in each inoculum for any ratio, except for the lung samples from hamsters 2, 10, and 19 (**Fig. 4**). For animals inoculated with a 1:1 or 1:3 ratio of BA.2.75:BA.5, the lung and nasal turbinate samples showed a greater proportion of BA.2.75, except for the lung sample from hamsters 2, 9, and 10 (**Fig. 4a and 4b**). Of note, even though the proportion of BA.2.75 in the inoculum was much lower than that of BA.5 (i.e., a 1:19 or 1:199 mixture of BA.2.75:BA.5), BA.2.75 became dominant in the lungs of four (#s 11, 12, 15, and 20) of the ten animals (**Fig. 4c and 4d**). Taken together, these results suggest that BA.2.75 may have greater replicative fitness than BA.5, especially in the upper respiratory tract.

**Figure 4.**
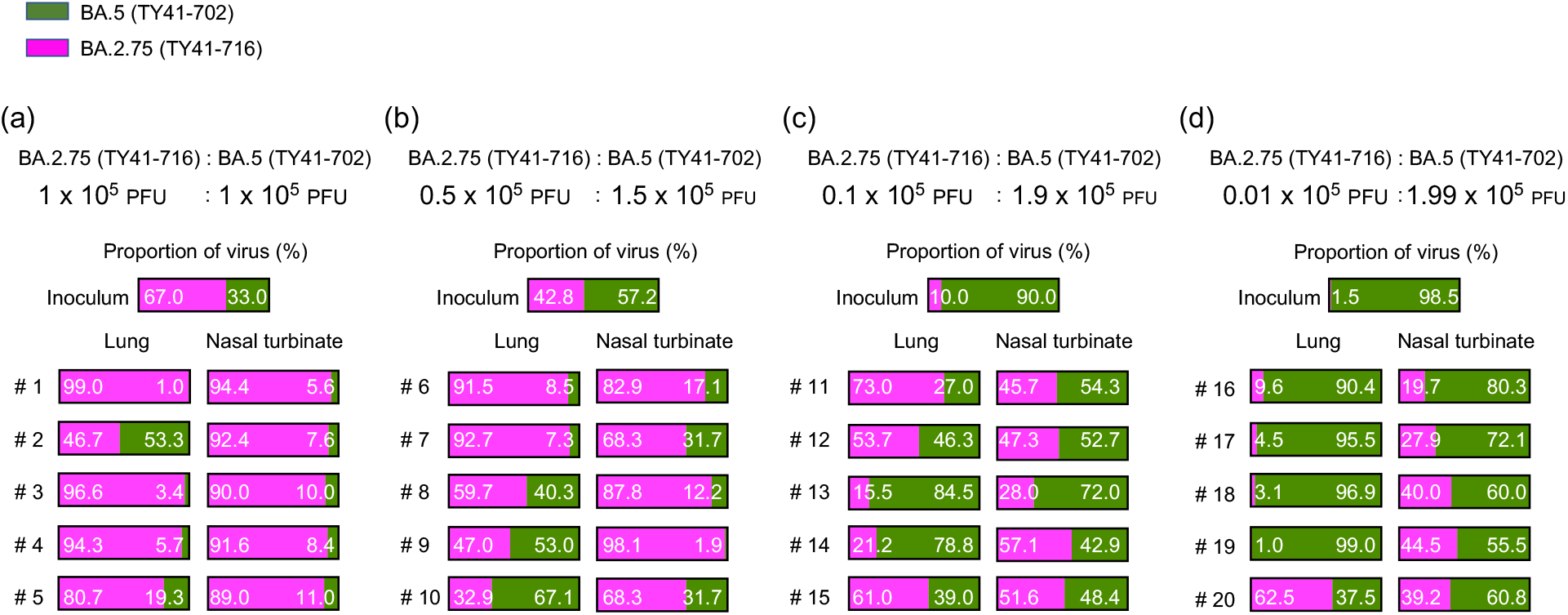
Relative viral fitness of BA.2.75 and BA.5 in hamsters. BA.2.75 (TY41-716) and BA.5 (TY41-702) were mixed at a 1:1 (**a**), 1:3 (**b**), 1:19 (**c**), or 1:199 (**d**) ratio on the basis of their infectious titers, and the virus mixture (total 2 × 10^5^ PFU in 60 µL) was intranasally inoculated into wild-type hamsters (*n* =5). Nasal turbinates and lungs were collected from the infected animals at 4 dpi and analyzed using next generation sequencing (NGS). Shown are the relative proportions of BA.5 and BA.2.75 in the infected animals.

## Discussion

We previously showed that Omicron sublineage BA.2 and BA.5 variants exhibit similar pathogenicity in rodent models by using several clinical isolates, and showed that both variants are significantly attenuated compared to previous circulating VOCs (Kawaoka et al., 2022; Uraki et al., 2022a). Here, we evaluated the replication and pathogenicity of Omicron sublineage BA.2.75 variants in hamsters. Our data show that there are no substantial differences in weight change among hamsters infected with BA.2.75, BA.2, or BA.5 (**Fig. 2a**); however, viral titers in the lungs of BA.2.75-infected hamsters were higher than those in the lungs of BA.2- or BA.5-infected hamsters (**Fig. 2c**). In addition, in competition assays, BA.2.75 replicated better than BA.5 in the lungs (**Fig. 4**). Of note, in the lungs of BA.2.75-inoculated hamsters, we observed focal pneumonia, characterized by patchy inflammation interspersed in alveolar regions, indicating that BA.2.75 can cause mild pneumonia (**Fig. 3 and S5**). In contrast, BA.5 mainly affected the bronchi, resulting in bronchitis/bronchiolitis, and did not cause obvious pneumonia (**Fig. 3**). Similar results were observed with hamsters infected with BA.1, BA.2, or BA.4 ((Halfmann et al., 2022; Kawaoka et al., 2022; Uraki et al., 2022b)). These findings suggest that among the Omicron variants, the Omicron subvariant BA.2.75 causes the most severe tissue damage in the lungs of hamsters.

Omicron variants, including BA.1 or BA.2, are less likely than Delta variants to be associated with pneumonia in COVID-19 patients (Christensen et al., 2022; Kozlov, 2022; Li et al., 2022), consistent with our previous data obtained in a hamster model (Halfmann et al., 2022; Uraki et al., 2022a). However, in the present study, we found that BA.2.75 can cause focal viral pneumonia in hamsters, unlike the other Omicron variants (i.e., BA.1, BA.2, BA.4, and BA.5) (Halfmann et al., 2022; Kawaoka et al., 2022; Uraki et al., 2022a). The reason for this is unclear; however, it might be due to differences in the binding affinity of the S protein for hACE2 among BA.1, BA.2, BA.4, BA.5, and BA.2.75. Recent studies have reported that the RBD of BA.2.75 exhibits higher binding affinity for the hACE2 receptor than that of BA.2 and BA.4/5 (Cao et al., 2022; Saito et al., 2022). SARS-CoV-2 enters cells in two distinct ways: by fusion of the viral lipid envelope with the target cell plasma membrane or fusion of the viral envelope with the endosomal membrane after internalization through the endocytic pathway (Hoffmann et al., 2020; Jackson et al., 2022; Walls et al., 2020). The internalization of SARS-CoV-2 via the endocytic pathway is believed to be induced by the binding of the virus to ACE2 (Bayati et al., 2021; Inoue et al., 2007). The Omicron variants have been shown to preferentially utilize the endocytic pathway to enter cells (Hui et al., 2022; Meng et al., 2022). In addition, previous studies have demonstrated that the enhanced binding affinity between ACE2 and the RBD increases the efficiency of SARS-CoV-2 entry (Ou et al., 2021; Ozono et al., 2021). Therefore, the higher ACE2 binding affinity of BA.2.75 may enhance its ability to infect the lungs, thereby allowing BA.2.75 to cause viral pneumonia in hamsters. Also, this higher ACE2 affinity of BA.2.75 may increase its competitive fitness compared to BA.5 in the respiratory tracts of hamsters, as observed in our *in vivo* competition assay (**Fig. 4**). Further investigations are required to determine whether ACE2 binding affinity truly influences Omicron infection.

We note two key limitations in this study: (1) although hamsters are one of the most widely used animals that are known to be susceptible to SARS-CoV-2, including mice and non-human primates (Chan et al., 2020; Imai et al., 2020; Sia et al., 2020), it is unclear whether the BA.2.75 variant causes more clinically severe respiratory disease than other Omicron variants in humans; and (2) our study was performed in immunologically naïve animals; however, many people have already acquired immunity to SARS-CoV-2 through natural infection and/or vaccination. Therefore, it remains unclear whether our data reflect the clinical outcome in patients with immunity against SARS-CoV-2. Clinical studies are needed to corroborate our findings in the hamster model.

In summary, the prevalence of BA.2.75 has increased throughout India, and has been increasing faster in regions where BA.5 and BA.2.75 are co-circulating, suggesting the potential for BA.2.75 to become the next globally dominant variant. Our data show that, compared to BA.5 and BA.2, BA.2.75 can replicate efficiently in the lungs of hamsters and cause more severe respiratory disease. This higher replicative ability of BA.2.75 in the lower respiratory tract may affect the clinical outcome in infected humans. Accordingly, the spread of this new variant should be monitored closely.

## Materials and Methods

### Variant tracking strategies

Figures 1, S2, and S3 show transitions between variant forms, emphasizing the recent expansion of the BA.2.75 variant that is indicative of a possible selective advantage. Details of the methods used to make these figures are described in Korber et al. (Korber et al., 2020), and web-based updates of these figures based on recent GISAID data can be generated via the “Embers” and “Isotonic Regression” tools at the Los Alamos National Laboratory SARS-CoV-2 variant analysis website (https://cov.lanl.gov). Figure 1 was created using Embers and displays running weekly counts and proportions of variants at different geographic levels. The Isotonic Regression analysis explores the dynamics of the transition towards higher frequencies of BA.2.75 over time, testing whether it is increasing in frequency relative to other variant forms at the country or state level, everywhere globally that BA.2.75 is well enough established to have been sampled ten or more times. A resampling statistic was used to evaluate whether the increasing sampling of BA.2.75 is significant (Elbe and Buckland-Merrett, 2017; Khare et al., 2021; Korber et al., 2020). The data sets for these figures were uploaded from GISAID 2022-08-15 (Elbe and Buckland-Merrett, 2017; Khare et al., 2021).

### Cells

VeroE6/TMPRSS2 (JCRB 1819) cells (Matsuyama et al., 2020) were propagated in the presence of 1 mg/ml geneticin (G418; Invivogen) and 5 μg/ml plasmocin prophylactic (Invivogen) in Dulbecco’s modified Eagle’s medium (DMEM) containing 10% Fetal Calf Serum (FCS). Vero E6-TMPRSS2-T2A-ACE2 cells (provided by Dr. Barney Graham, NIAID Vaccine Research Center) were cultured in DMEM supplemented with 10% FCS, 10 mM HEPES pH 7.3, 100 U/mL penicillin–streptomycin, and 10 μg/mL puromycin. VeroE6/TMPRSS2 and Vero E6-TMPRSS2-T2A-ACE2 cells were maintained at 37 L with 5% CO_2_. The cells were regularly tested for mycoplasma contamination by using PCR, and confirmed to be mycoplasma-free.

### Viruses

hCoV-19/Japan/UT-NCD1288-2N/2022 (BA.2; NCD1288, Accession ID; EPI_ISL_9595604) (Takashita et al., 2022; Uraki et al., 2022a), hCoV-19/Japan/TY41-716/2022 (BA.2.75; TY41-716, Accession ID; EPI_ISL_14011362)(Takashita et al., 2022), hCoV-19/Japan/UT-NCD1757-1N/2022 (BA.2.75; NCD1757, Accession ID; EPI_ISL_14321758), hCoV-19/Japan/UT-NCD1759-1N/2022 (BA.2.75; NCD1759, Accession ID; EPI_ISL_14321760), hCoV-19/Japan/TY41-702/2022 (BA.5; TY41-702, Accession ID; EPI_ISL_13512581) (Kawaoka et al., 2022), and hCoV-19/USA/WI-UW-5250/2021 (B.1.617.2; UW5250) (Gagne et al., 2022; Halfmann et al., 2022) were propagated in VeroE6/TMPRSS2 cells in VP-SFM (Thermo Fisher Scientific). BA.2.75 (NCD1757) and BA.2.75 (NCD1759) were subjected to next generation sequencing (NGS) (see Whole genome sequencing). All experiments with SARS-CoV-2 were performed in enhanced biosafety level 3 (BSL3) containment laboratories at the University of Tokyo and the National Institute of Infectious Diseases, Japan, which are approved for such use by the Ministry of Agriculture, Forestry, and Fisheries, Japan, or in BSL3 agriculture containment laboratories at the University of Wisconsin-Madison, which are approved for such use by the Centers for Disease Control and Prevention and by the US Department of Agriculture.

### Animal experiments and approvals

Animal studies were carried out in accordance with the recommendations in the Guide for the Care and Use of Laboratory Animals of the National Institutes of Health. The protocols were approved by the Animal Experiment Committee of the Institute of Medical Science, the University of Tokyo (approval number PA19-75) and the Institutional Animal Care and Use Committee at the University of Wisconsin, Madison (assurance number V006426). Virus inoculations were performed under isoflurane, and all efforts were made to minimize animal suffering. *In vivo* studies were not blinded, and animals were randomly assigned to infection groups. No sample-size calculations were performed to power each study. Instead, sample sizes were determined based on prior *in vivo* virus challenge experiments.

### Experimental infection of Syrian hamsters

Six-week-old male wild-type Syrian hamsters (Japan SLC Inc., Shizuoka, Japan) were used in this study. Baseline body weights were measured before infection. Under isoflurane anesthesia, five hamsters per group were intranasally inoculated with 10^5^ PFU (in 30 μL) of BA.2 (NCD1288), BA.2.75 (TY41-716), BA.2.75 (NCD1757), BA.2.75 (NCD1759), BA.5 (TY41-702), or B.1.617.2 (UW5250). Body weight was monitored daily for 6 days. For virological and pathological examinations, ten hamsters per group were intranasally infected with 10^5^ PFU (in 30 μL) of BA.2 (NCD1288), BA.2.75 (TY41-716), BA.2.75 (NCD1757), BA.5(TY41-702), or B.1.617.2 (UW5250); 3 and 6 dpi, five animals were euthanized and nasal turbinates and lungs were collected. The virus titers in the nasal turbinates and lungs were determined by use of plaque assays on Vero E6-TMPRSS2-T2A-ACE2 cells.

For co-infection studies, BA.2.75 (TY41-716) was mixed with BA.5 (TY41-702) at a 1:1, 1:3, 1:19, or 1:199 ratio on the basis of their titers, and each virus mixture (total 2 × 10^5^ PFU in 60 µL) was inoculated into five wild-type hamsters. At 4 dpi, five animals were euthanized and nasal turbinates and lungs were collected to determine virus titers.

The K18-hACE2 transgenic hamster line (line M41) were developed by using a piggyBac-mediated transgenic approach. The K18-hACE2 cassette from the pK18-hACE2 plasmid was transferred into a piggyBac vector, pmhyGENIE-3, for pronuclear injection (Gilliland et al., 2021). Then, female 6-8-week-old K18-hACE2 homozygous transgenic hamsters, whose hACE2 expression was confirmed, were intranasally inoculated with 10^5^ PFU (in 30 μL) of BA.2 (NCD1288), BA.5 (TY41-702), BA.2.75 (TY41-716), or B.1.617.2 (UW5250). At 5 dpi, the animals were euthanized and nasal turbinates and lungs were collected. The virus titers in the nasal turbinates and lungs were determined by use of plaque assays on Vero E6-TMPRSS2-T2A-ACE2 cells.

### Lung function

Respiratory parameters were measured by using a whole-body plethysmography system (PrimeBioscience) according to the manufacturer’s instructions. In brief, infected hamsters were placed in the unrestrained plethysmography chambers and allowed to acclimate for 1 min before data were acquired over a 3-min period by using FinePointe software.

### Histopathology

Histopathological examination was performed as previously described (Halfmann et al., 2022; Kawaoka et al., 2022; Uraki et al., 2022a). In brief, excised animal lungs were fixed in 4% paraformaldehyde in phosphate buffered saline (PBS) and processed for paraffin embedding. The paraffin blocks were sliced into 3µm-thick sections and mounted on silane-coated glass slides, followed by hematoxylin and eosin (H&E) stain for histopathological examination. To detect SARS-CoV-2 RNA, in situ hybridization was performed using an RNA scope 2.5 HD Red Detection kit (Advanced Cell Diagnostics, Newark, California) with an antisense probe targeting the nucleocapsid gene of SARS-CoV-2 (Advanced Cell Diagnostics) following manufacturer’s instructions. Tissue sections were also processed for immunohistochemistry with a rabbit polyclonal antibody for SARS-CoV nucleocapsid protein (ProSpec; ANT-180, 1:500 dilution, Rehovot, Israel), which cross-reacts with SARS-CoV-2 nucleocapsid protein. Specific antigen-antibody reactions were visualized by means of 3,3’-diaminobenzidine tetrahydrochloride staining using the Dako Envision system (Dako Cytomation; K4001, 1:1 dilution, Glostrup, Denmark).

### Whole genome sequencing

Viral RNA was extracted by using a QIAamp Viral RNA Mini Kit (QIAGEN). The whole genome of SARS-CoV-2 was amplified by using a modified ARTIC network protocol in which some primers were replaced or added (Itokawa et al., 2020; Quick). Briefly, viral cDNA was synthesized from the extracted RNA by using a LunarScript RT SuperMix Kit (New England BioLabs). The DNA was then amplified by performing a multiplexed PCR in two pools using the ARTIC-N5 primers and the Q5 Hot Start DNA polymerase (New England BioLabs) (Itokawa et al.). The DNA libraries for Illumina NGS were prepared from pooled amplicons by using a QIAseq FX DNA Library Kit (QIAGEN) and were then analyzed by using the iSeq 100 System (Illumina). To determine the sequences of BA.2.75 (NCD1757) and BA.2.75 (NCD1759), the reads were assembled by the CLC Genomics Workbench (version 22, Qiagen) with the Wuhan/Hu-1/2019 sequence (GenBank accession no. MN908947) as a reference. The sequences of BA.2.75 (NCD1757) and BA.2.75 (NCD1759) were deposited in the Global Initiative on Sharing All Influenza Data (GISAID) database with Accession IDs: EPI_ISL_14321758, and EPI_ISL_14321760, respectively. For the analysis of the ratio of BA.5 to BA.2.75 after co-infection, the ratio of BA.2.75 to BA.5 was calculated from the 6 amino acid differences in the S gene between the two viruses. Samples with more than 300 read-depths were analyzed.

### Statistical analysis

GraphPad Prism software was used to analyze the data. Statistical analysis included the Kruskal-Wallis test followed by Dunn’s test, and an ANOVA with post-hoc tests. Differences among groups were considered significant for *P* values < 0.05.

### Data and code availability

All data supporting the findings of this study are available in the paper. There are no restrictions in obtaining access to the primary data. The source data are provided with this paper. No novel code was used in the course of the data acquisition or analysis, the most representative forms of each virus, dynamics plots, and isotonic regression analyses are available at https://cov.lanl.gov/content/index.

## Acknowledgements

We thank Susan Watson for scientific editing. We also thank Kyoko Yokota, Naoko Mizutani, Kengo Kajiyama, Yuko Sato, and Seiya Ozono for technical assistance. We thank Hyejin Yoon for ongoing work to maintain cov.lanl.gov, and the development team at GISAID for supporting our Los Alamos effort. Vero E6-TMPRSS2-T2A-ACE2 cells were provided by Dr. Barney Graham, NIAID Vaccine Research Center. This work was supported by a Research Program on Emerging and Re-emerging Infectious Diseases (JP21fk0108615, JP21fk0108552 and JP22fk0108637), the Japan Program for Infectious Diseases Research and Infrastructure (JP22wm0125002, JP22wm0125008) from the Japan Agency for Medical Research and Development (AMED), the National Institutes of Health SARS-CoV-2 Assessment of Viral Evolution (SAVE) Project (AAI22018-001), the National Institutes of Allergy and Infectious Diseases Center for Research on Influenza Pathogenesis (HHSN272201400008C), and the Center for Research on Influenza Pathogenesis and Transmission (CRIPT) (75N93021C00014).

## Author contributions

R.U., S.I., P.J.H, S.Y., Y.H., K.I.-H., M.Kiso, M.Ito, Y.F., H.U., S.M., M.Kuroda, T.M., T.K., S.M., M.Imai, and T.S performed the hamster infection experiments, titrated virus in tissues, and/or analyzed pathology. S.Y. performed next generation sequencing. Z.W., R.L., Y.L., and D.L. generated human ACE2 hamsters. S.Y., Y.S.-T., N.I., S.F., S.W., K.M., and N.O. propagated and/or sequenced viruses. J.T. and B.K. analyzed variant dynamics and Spike genomes. W. F. processed viral sequence data for identifying variant-representative full-length genomes. R.U., S.I., P.J.H, S.Y., M.Imai, T.S. and Y.K. obtained funding, conceived the study, and/or supervised the research. R.U., M.Imai and Y.K. wrote the initial draft, with all other authors providing editorial comments.

## Corresponding author

Correspondence to: Yoshihiro Kawaoka, D.V.M., Ph.D.; Tadaki Suzuki, M.D., Ph.D.; Masaki Imai, D.V.M., Ph.D.; and Bette Korber, Ph.D..

## Figure legends

**Figure S1. Amino acid differences between representative forms of recently emerged Omicron variants**.

**a**, Amino acid differences in the Spike of Omicron variants. BA.1 was rapidly globally replaced by BA.2; here we used the most common form of BA.2 as the reference Omicron variant. Spike amino acid differences between the Wuhan reference strain WIV04/2019|EPI_ISL_402124|2019 and the baseline form of BA.2 are shown in black. When other Omicron variants share spike BA.2 defining mutations in a given position, they are noted in grey. When they differ, the amino acid change is highlighted in the color assigned to each variant (the same color as used in Figure 1). Deletions are indicated by a dash (-), insertions by a plus sign (e.g., +214EPE means a three amino acid insertion of EPE after position 214). Reversions from BA.2 to the ancestral Wuhan form are indicated by an underscore (_). **b**, Highlighting amino acid differences between BA.2, BA.5, and BA.2.75 throughout the full proteome. Only amino differences from the most representative form of BA.2 are shown in this figure, illustrated as a tick mark. The grey line represents the full proteome. All changes in the most common forms of BA.5 and BA.2.75 relative to BA.2 are noted, as these are candidates for contributing to a selective advantage of BA.5 over BA.2, and of BA.2.75 over BA.5 and BA.2. Because details for the spike protein are shown in **a**, they are not shown here. The amino acids that are distinctive in the three BA.2.75 variants studied in this paper are highlighted at the bottom, BA.2.75 V1–V3. Full length representative forms of Pango lineages are defined as the most common circulating form of a given Pango lineage.

**Figure S2. Isotonic regression analysis showing BA.2.75 is increasingly sampled over time in countries where it has become established**.

The table provides summary statistics for all countries where BA.2.75 sequences have been sampled more than 10 times with a sampling date between 12-05-2022 and 08-08-2022. Four examples of the data over time are plotted to illustrate the increasing frequency of BA.2.75 sampling. We show the countries with the most available data (India, Singapore, and the US) as well as Nepal, with a lower sampling frequency but higher fraction of BA.2.75 cases – these numbers are highlighted in red in the table. The proportion of BA.27.5 in the total sample (y-axis) is calculated each day samples are available (x-axis). The size of the dot reflects the relative sample size on a given day. The *p*-value is calculated based on a one-sided resampling test with 400 randomizations. All *p*-values in the right-hand column are highly significant, showing that the frequency of BA.2.75 is increasing in every country where it has been sampled more than ten times. The results can be updated using the Isotonic Regression tool at cov.lanl.gov. (https://cov.lanl.gov/content/sequence/ISORG/pango_isorg.html)(Korber et al., 2020). An additional analysis was performed comparing BA.2.75 to just BA.5 at the country level, and these direct comparisons also supported the conclusion that BA.2.75 is expanding significantly faster than BA.5 in regions where they are co-circulating.

**Figure S3. Isotonic regression analysis showing BA.2.75 is increasingly sampled over time in all states in India where it has been sampled more than 10 times**.

These figures follow the format of Figure S2, but at a more geographically restricted. The analysis establishes that BA.2.75 is increasing in frequency throughout India, providing evidence that the increase at the country level in India seen in Figs. 1 and S2 are not due to regional founder effects within India. The four Indian states with the highest number of samples (the values in red) were selected to illustrate the increases in the lower panels. BA.2.75 prevalence is also increasing in the four US states and two Canadian provinces where it has been sampled more than 10 times, as well as in England and New South Wales, Australia. An additional analysis was performed where we compared BA.2.75 to *just* BA.5 at the state level, and these direct comparisons also supported the conclusion that BA.2.75 is expanding significantly faster than BA.5 in regions where they are co-circulating.

**Figure S4. Semi-macroscopic images of the lungs of hamsters inoculated with SARS-CoV-2**.

Syrian hamsters (n = 5, per group) were inoculated with 10^5^ PFU of BA.2.75 (TY41-716), BA.5 (TY41-702), or B.1.617.2 (UW5250) and sacrificed at 3 or 6 dpi for histopathological examinations. Semi-macroscopic images (hematoxylin and eosin staining) of the lungs from all animals examined are shown. Scale bars, 5 mm.

**Figure S5. Histopathological findings in hamsters inoculated with BA.2.75 (NCD1757)**.

Syrian hamsters (n = 5) were inoculated with 10^5^ PFU of BA.2.75 (Omicron, NCD1757) and sacrificed at 6 dpi for histopathological examinations. **a**, Semi-macroscopic images (hematoxylin and eosin staining) of the lungs from all animals examined are shown. Scale bars, 5 mm. **b**, Representative images (hematoxylin and eosin staining) of the lungs at low magnification are shown. Scale bars, 1 mm. **c**, Representative images (hematoxylin and eosin staining) of the bronchi/bronchioles and alveoli at high magnification are shown. Scale bars, 100 µm.

## References

Bayati, A., Kumar, R., Francis, V., and McPherson, P.S. (2021). SARS-CoV-2 infects cells after viral entry via clathrin-mediated endocytosis. J Biol Chem. 296, 100306. Published online 2021/01/22 DOI: 10.1016/j.jbc.2021.100306.

Cao, Y., Yu, Y., Song, W., Jian, F., Yisimayi, A., Yue, C., Feng, R., Wang, P., Yu, L., Zhang, N., et al. (2022). Neutralizing antibody evasion and receptor binding features of SARS-CoV-2 Omicron BA.2.75. bioRxiv. DOI: 10.1101/2022.07.18.500332.

Chan, J.F., Zhang, A.J., Yuan, S., Poon, V.K., Chan, C.C., Lee, A.C., Chan, W.M., Fan, Z., Tsoi, H.W., Wen, L., et al. (2020). Simulation of the Clinical and Pathological Manifestations of Coronavirus Disease 2019 (COVID-19) in a Golden Syrian Hamster Model: Implications for Disease Pathogenesis and Transmissibility. Clin Infect Dis. 71(9), 2428–2446. Published online 2020/03/28 DOI: 10.1093/cid/ciaa325.

Christensen, P.A., Olsen, R.J., Long, S.W., Snehal, R., Davis, J.J., Ojeda Saavedra, M., Reppond, K., Shyer, M.N., Cambric, J., Gadd, R., et al. (2022). Signals of Significantly Increased Vaccine Breakthrough, Decreased Hospitalization Rates, and Less Severe Disease in Patients with Coronavirus Disease 2019 Caused by the Omicron Variant of Severe Acute Respiratory Syndrome Coronavirus 2 in Houston, Texas. Am J Pathol. 192(4), 642–652. Published online 2022/02/07 DOI: 10.1016/j.ajpath.2022.01.007.

Elbe, S., and Buckland-Merrett, G. (2017). Data, disease and diplomacy: GISAID’s innovative contribution to global health. Glob Chall. 1(1), 33–46. Published online 2017/01/10 DOI: 10.1002/gch2.1018.

Flemming, A. (2022). Omicron, the great escape artist. Nat Rev Immunol. 22(2), 75. Published online 2022/01/13 DOI: 10.1038/s41577-022-00676-6.

Gagne, M., Corbett, K.S., Flynn, B.J., Foulds, K.E., Wagner, D.A., Andrew, S.F., Todd, J.M., Honeycutt, C.C., McCormick, L., Nurmukhambetova, S.T., et al. (2022). Protection from SARS-CoV-2 Delta one year after mRNA-1273 vaccination in rhesus macaques coincides with anamnestic antibody response in the lung. Cell. 185(1), 113–130 e115. Published online 2021/12/19 DOI: 10.1016/j.cell.2021.12.002.

Gilliland, T., Liu, Y., Li, R., Dunn, M., Cottle, E., Terada, Y., Ryckman, Z., Alcorn, M., Vasilatos, S., Lundy, J., et al. (2021). Protection of human ACE2 transgenic Syrian hamsters from SARS CoV-2 variants by human polyclonal IgG from hyper-immunized transchromosomic bovines. bioRxiv. Published online 2021/08/04 DOI: 10.1101/2021.07.26.453840.

Halfmann, P.J., Iida, S., Iwatsuki-Horimoto, K., Maemura, T., Kiso, M., Scheaffer, S.M., Darling, T.L., Joshi, A., Loeber, S., Singh, G., et al. (2022). SARS-CoV-2 Omicron virus causes attenuated disease in mice and hamsters. Nature. Published online 2022/01/22 DOI: 10.1038/s41586-022-04441-6.

Hoffmann, M., Kleine-Weber, H., Schroeder, S., Kruger, N., Herrler, T., Erichsen, S., Schiergens, T.S., Herrler, G., Wu, N.H., Nitsche, A., et al. (2020). SARS-CoV-2 Cell Entry Depends on ACE2 and TMPRSS2 and Is Blocked by a Clinically Proven Protease Inhibitor. Cell. 181(2), 271–280 e278. Published online 2020/03/07 DOI: 10.1016/j.cell.2020.02.052.

Hui, K.P.Y., Ho, J.C.W., Cheung, M.C., Ng, K.C., Ching, R.H.H., Lai, K.L., Kam, T.T., Gu, H., Sit, K.Y., Hsin, M.K.Y., et al. (2022). SARS-CoV-2 Omicron variant replication in human bronchus and lung ex vivo. Nature. 603(7902), 715–720. Published online 2022/02/02 DOI: 10.1038/s41586-022-04479-6.

Imai, M., Iwatsuki-Horimoto, K., Hatta, M., Loeber, S., Halfmann, P.J., Nakajima, N., Watanabe, T., Ujie, M., Takahashi, K., Ito, M., et al. (2020). Syrian hamsters as a small animal model for SARS-CoV-2 infection and countermeasure development. Proc Natl Acad Sci U S A. 117(28), 16587–16595. Published online 2020/06/24 DOI: 10.1073/pnas.2009799117.

Inoue, Y., Tanaka, N., Tanaka, Y., Inoue, S., Morita, K., Zhuang, M., Hattori, T., and Sugamura, K. (2007). Clathrin-dependent entry of severe acute respiratory syndrome coronavirus into target cells expressing ACE2 with the cytoplasmic tail deleted. J Virol. 81(16), 8722–8729. Published online 2007/05/25 DOI: 10.1128/JVI.00253-07.

Itokawa, K., Sekizuka, T., Hashino, M., and al., e., nCoV-2019 sequencing protocol for illumina V.5 Itokawa, K., Sekizuka, T., Hashino, M., Tanaka, R., and Kuroda, M. (2020). Disentangling primer interactions improves SARS-CoV-2 genome sequencing by multiplex tiling PCR. PLoS One. 15(9), e0239403. Published online 2020/09/19 DOI: 10.1371/journal.pone.0239403.

Jackson, C.B., Farzan, M., Chen, B., and Choe, H. (2022). Mechanisms of SARS-CoV-2 entry into cells. Nat Rev Mol Cell Biol. 23(1), 3–20. Published online 2021/10/07 DOI: 10.1038/s41580-021-00418-x.

Kawaoka, Y., Uraki, R., Halfmann, P., Iida, S., Yamayoshi, S., Furusawa, Y., Kiso, M., Ito, M., Iwatsuki-Horimoto, K., Mine, S., et al. (2022). Characterization of SARS-CoV-2 Omicron BA.4 and BA.5 clinical isolates. Research Square. DOI: 10.21203/rs.3.rs-1820048/v1.

Khare, S., Gurry, C., Freitas, L., Schultz, M.B., Bach, G., Diallo, A., Akite, N., Ho, J., Lee, R.T., Yeo, W., et al. (2021). GISAID’s Role in Pandemic Response. China CDC Wkly. 3(49), 1049–1051. Published online 2021/12/23 DOI: 10.46234/ccdcw2021.255.

Korber, B., Fischer, W.M., Gnanakaran, S., Yoon, H., Theiler, J., Abfalterer, W., Hengartner, N., Giorgi, E.E., Bhattacharya, T., Foley, B., et al. (2020). Tracking Changes in SARS-CoV-2 Spike: Evidence that D614G Increases Infectivity of the COVID-19 Virus. Cell. 182(4), 812–827 e819. Published online 2020/07/23 DOI: 10.1016/j.cell.2020.06.043.

Kozlov, M. (2022). Omicron’s feeble attack on the lungs could make it less dangerous. Nature. 601(7892), 177. Published online 2022/01/07 DOI: 10.1038/d41586-022-00007-8.

Li, X., Wu, L., Qu, Y., Cao, M., Feng, J., Huang, H., Liu, Y., Lu, H., Liu, Q., and Liu, Y. (2022). Clinical characteristics and vaccine effectiveness against SARS-CoV-2 Omicron subvariant BA.2 in the children. Signal Transduct Target Ther. 7(1), 203. Published online 2022/06/29 DOI: 10.1038/s41392-022-01023-w.

Matsuyama, S., Nao, N., Shirato, K., Kawase, M., Saito, S., Takayama, I., Nagata, N., Sekizuka, T., Katoh, H., Kato, F., et al. (2020). Enhanced isolation of SARS-CoV-2 by TMPRSS2-expressing cells. Proc Natl Acad Sci U S A. 117(13), 7001–7003. Published online 2020/03/14 DOI: 10.1073/pnas.2002589117.

Meng, B., Abdullahi, A., Ferreira, I., Goonawardane, N., Saito, A., Kimura, I., Yamasoba, D., Gerber, P.P., Fatihi, S., Rathore, S., et al. (2022). Altered TMPRSS2 usage by SARS-CoV-2 Omicron impacts infectivity and fusogenicity. Nature. 603(7902), 706–714. Published online 2022/02/02 DOI: 10.1038/s41586-022-04474-x.

Ou, J., Zhou, Z., Dai, R., Zhang, J., Zhao, S., Wu, X., Lan, W., Ren, Y., Cui, L., Lan, Q., et al. (2021). V367F Mutation in SARS-CoV-2 Spike RBD Emerging during the Early Transmission Phase Enhances Viral Infectivity through Increased Human ACE2 Receptor Binding Affinity. J Virol. 95(16), e0061721. Published online 2021/06/10 DOI: 10.1128/JVI.00617-21.

Ozono, S., Zhang, Y., Ode, H., Sano, K., Tan, T.S., Imai, K., Miyoshi, K., Kishigami, S., Ueno, T., Iwatani, Y., et al. (2021). SARS-CoV-2 D614G spike mutation increases entry efficiency with enhanced ACE2-binding affinity. Nat Commun. 12(1), 848. Published online 2021/02/10 DOI: 10.1038/s41467-021-21118-2.

Quick, J., nCoV-2019 sequencing protocol. Saito, A., Tamura, T., Zahradnik, J., Deguchi, S., Tabata, K., Kimura, I., Ito, J., Nasser, H., Toyoda, M., Nagata, K., et al. (2022). Virological characteristics of the SARS-CoV-2 Omicron BA.2.75. bioRxiv. DOI: 10.1101/2022.08.07.503115.

Sia, S.F., Yan, L.M., Chin, A.W.H., Fung, K., Choy, K.T., Wong, A.Y.L., Kaewpreedee, P., Perera, R., Poon, L.L.M., Nicholls, J.M., et al. (2020). Pathogenesis and transmission of SARS-CoV-2 in golden hamsters. Nature. 583(7818), 834–838. Published online 2020/05/15 DOI: 10.1038/s41586-020-2342-5.

Takashita, E., Kinoshita, N., Yamayoshi, S., Sakai-Tagawa, Y., Fujisaki, S., Ito, M., Iwatsuki-Horimoto, K., Halfmann, P., Watanabe, S., Maeda, K., et al. (2022). Efficacy of Antiviral Agents against the SARS-CoV-2 Omicron Subvariant BA.2. N Engl J Med. 386(15), 1475–1477. Published online 2022/03/10 DOI: 10.1056/NEJMc2201933.

Takashita, E., Yamayoshi, S., Fukushi, S., Suzuki, T., Maeda, K., Sakai-Tagawa, Y., Ito, M., Uraki, R., Halfmann, P., Watanabe, S., et al. (2022). Efficacy of Antiviral Agents against the Omicron Subvariant BA.2.75. New England Journal of Medicine. in press.

Uraki, R., Kiso, M., Iida, S., Imai, M., Takashita, E., Kuroda, M., Halfmann, P.J., Loeber, S., Maemura, T., Yamayoshi, S., et al. (2022a). Characterization and antiviral susceptibility of SARS-CoV-2 Omicron BA.2. Nature. 607(7917), 119–127. Published online 2022/05/17 DOI: 10.1038/s41586-022-04856-1.

Uraki, R., Kiso, M., Imai, M., Yamayoshi, S., Ito, M., and Michiko Ujie, Y.F., Kiyoko Iwatsuki-Horimoto, Yuko Sakai-Tagawa, Yoshihiro Kawaoka (2022b). Therapeutic efficacy of antibodies and antivirals against a SARS-CoV-2 Omicron variant. Research Square. DOI: 10.21203/rs.3.rs-1240227/v1.

Walls, A.C., Park, Y.J., Tortorici, M.A., Wall, A., McGuire, A.T., and Veesler, D. (2020). Structure, Function, and Antigenicity of the SARS-CoV-2 Spike Glycoprotein. Cell. 181(2), 281–292 e286. Published online 2020/03/11 DOI: 10.1016/j.cell.2020.02.058.

