## Supplemental Figures for "Characterization of SARS-CoV-2 Omicron BA.2.75 clinical isolates"

### Supplementary Fig. 1

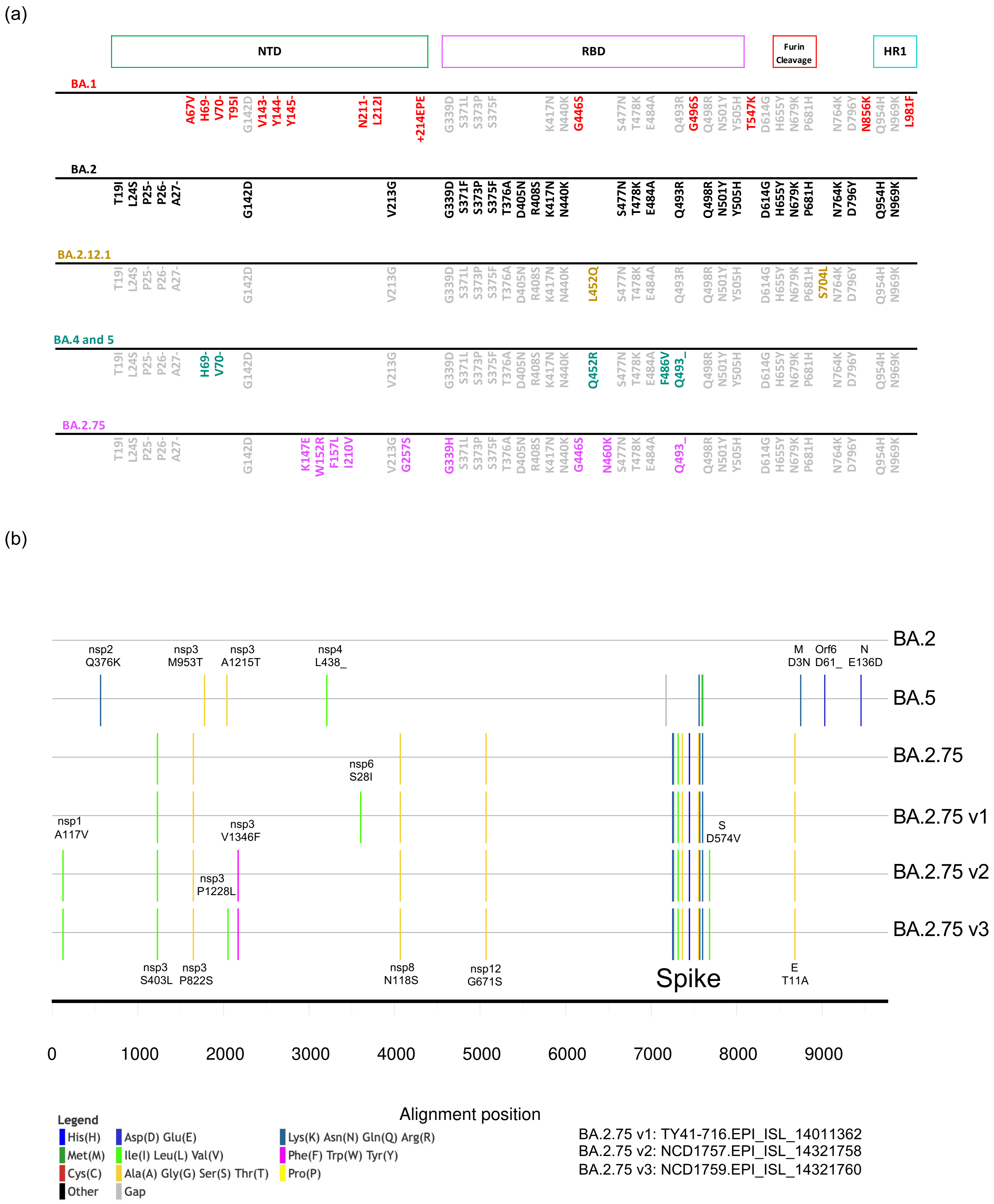

### Supplementary Fig.2

| | Countries where BA.2.75 has been sampled ≥10 times | Number of BA.2.75 sequences | Number of other sequences | Total Number of sequences | BA.2.75/Total (%) | Number of days | Time window | One-sided *P*-value, increasing |
|---|---|---|---|---|---|---|---|---|
| **Country level** | Australia | 33 | 26146 | 26179 | 0.13 | 83 | 82 | 0.00249 |
| | Canada | 33 | 33414 | 33447 | 0.1 | 77 | 76 | 0.00249 |
| | Denmark | 13 | 40112 | 40125 | 0.03 | 81 | 80 | 0.00249 |
| | India | 994 | 12717 | 13711 | 7.25 | 78 | 77 | 0.00249 |
| | Israel | 11 | 37764 | 37775 | 0.03 | 80 | 79 | 0.00249 |
| | Japan | 29 | 38221 | 38250 | 0.08 | 81 | 80 | 0.00249 |
| | Nepal | 17 | 136 | 153 | 11.11 | 48 | 66 | 0.00249 |
| | Singapore | 42 | 4135 | 4177 | 1.01 | 77 | 76 | 0.00249 |
| | USA | 82 | 365878 | 365960 | 0.02 | 83 | 82 | 0.00249 |
| | United-Kingdom | 32 | 89827 | 89859 | 0.04 | 82 | 81 | 0.00249 |

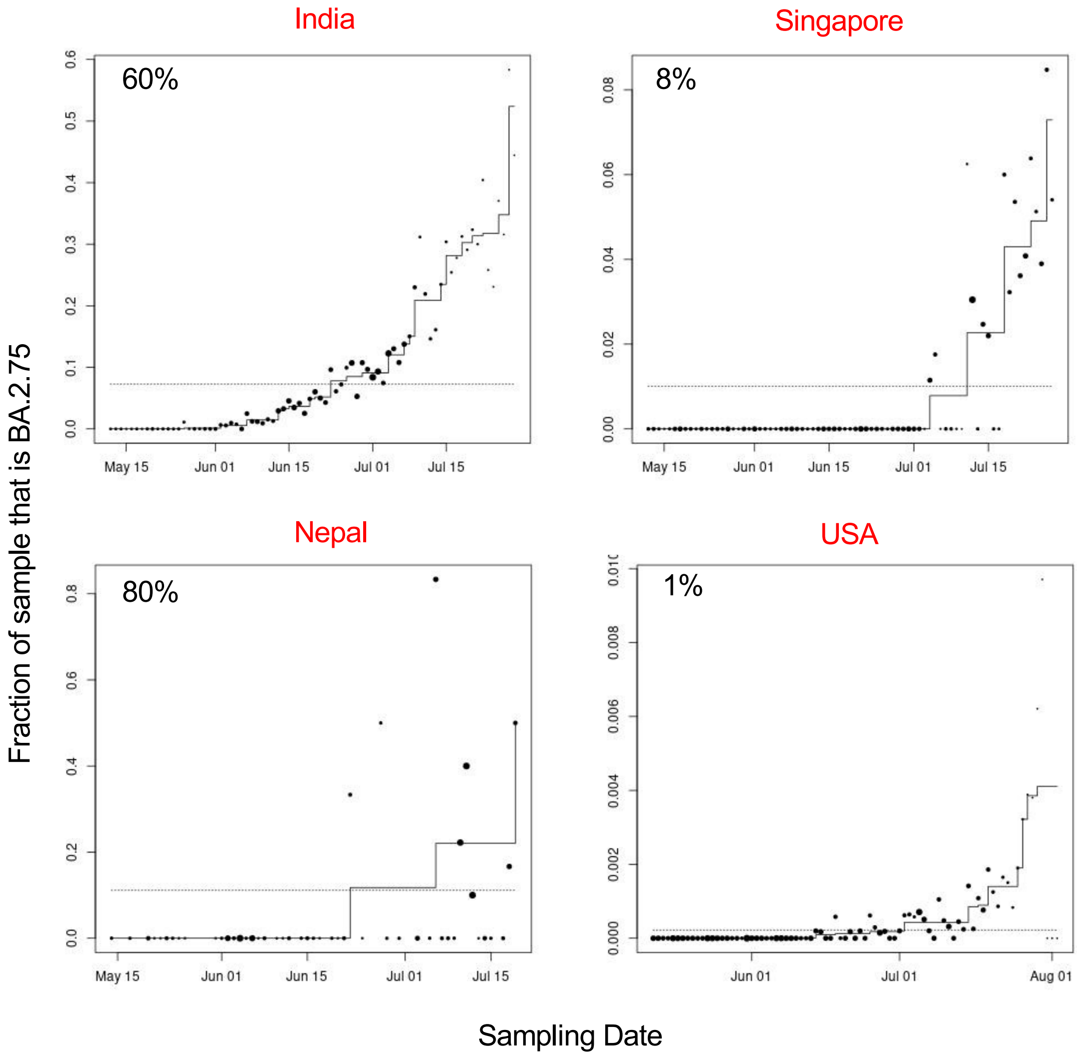

### Supplementary Fig. 3

| | Countries where BA.2.75 has been sampled ≥10 times | Number of BA.2.75 sequences | Number of other sequences | Total Number of sequences | BA.2.75/Total (%) | Number of days | Time window | One-sided *P*-value, increasing |
|---|---|---|---|---|---|---|---|---|
| State Level | Australia_New-South-Wales | 21 | 6961 | 6982 | 0.3 | 83 | 82 | 0.00249 |
| | Canada_Alberta | 10 | 5090 | 5100 | 0.2 | 72 | 71 | 0.01244 |
| | Canada_Ontario | 21 | 13167 | 13188 | 0.16 | 76 | 75 | 0.00249 |
| | India_Assam | 20 | 338 | 358 | 5.59 | 28 | 49 | 0.00249 |
| | India_Chhattisgarh | 10 | 99 | 109 | 9.17 | 31 | 43 | 0.20647 |
| | India_Delhi | 71 | 1238 | 1309 | 5.42 | 71 | 75 | 0.00249 |
| | India_Gujarat | 44 | 810 | 854 | 5.15 | 60 | 65 | 0.00249 |
| | India_Haryana | 31 | 196 | 227 | 13.66 | 40 | 58 | 0.00249 |
| | India_Himachal-Pradesh | 37 | 113 | 150 | 24.67 | 51 | 64 | 0.00249 |
| | India_Maharashtra | 232 | 2532 | 2764 | 8.39 | 78 | 77 | 0.00249 |
| | India_Odisha | 135 | 407 | 542 | 24.91 | 61 | 74 | 0.00249 |
| | India_Rajasthan | 65 | 617 | 682 | 9.53 | 35 | 61 | 0.00249 |
| | India_Tamil-Nadu | 23 | 1913 | 1936 | 1.19 | 61 | 60 | 0.00249 |
| | India_Telangana | 73 | 1276 | 1349 | 5.41 | 76 | 75 | 0.00249 |
| | India_West-Bengal | 231 | 2576 | 2807 | 8.23 | 62 | 64 | 0.00249 |
| | USA_California | 16 | 65538 | 65554 | 0.02 | 80 | 79 | 0.00249 |
| | USA_New-Jersey | 10 | 15482 | 15492 | 0.06 | 77 | 76 | 0.00249 |
| | USA_New-York | 10 | 40566 | 40576 | 0.02 | 77 | 76 | 0.00249 |
| | USA_Washington | 10 | 14167 | 14177 | 0.07 | 74 | 74 | 0.00249 |
| | United-Kingdom_England | 26 | 66154 | 66180 | 0.04 | 82 | 81 | 0.00249 |

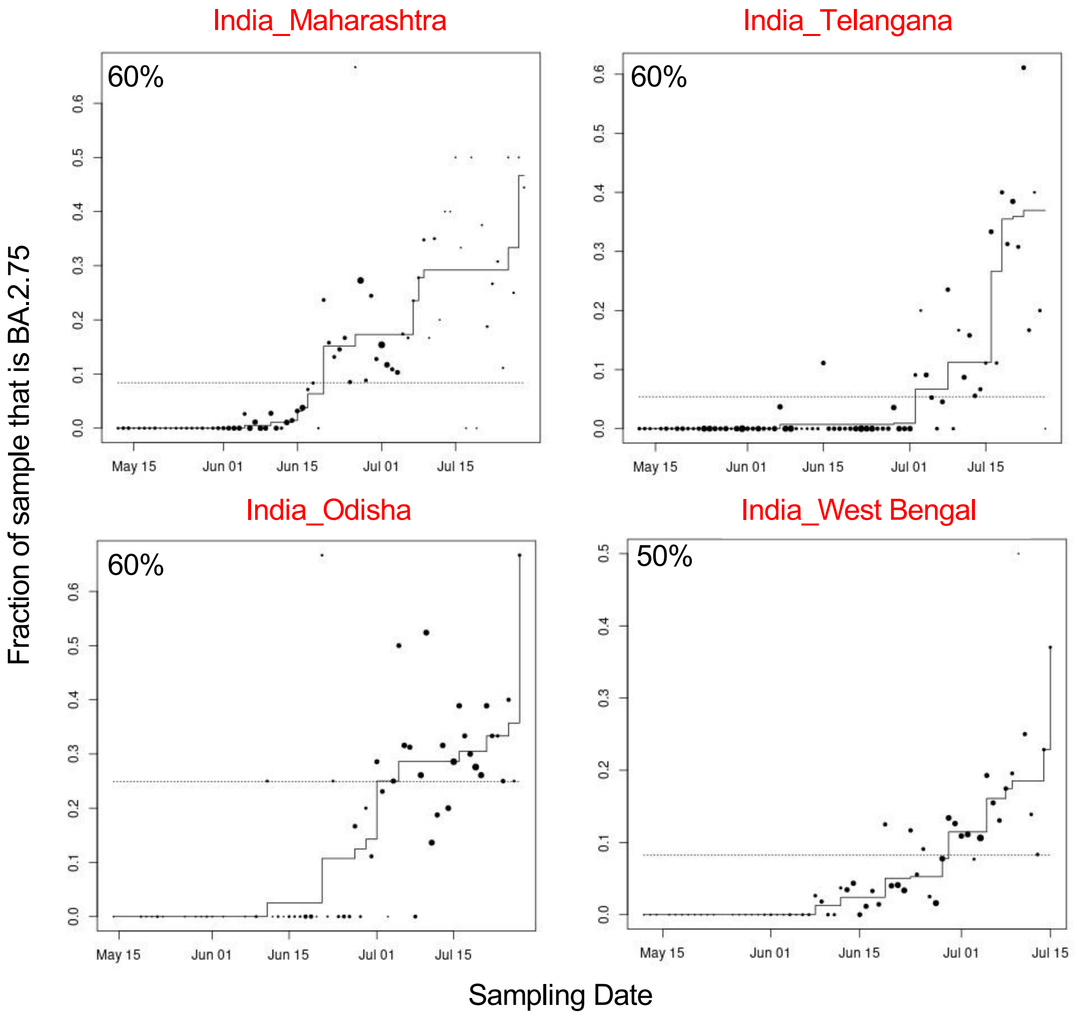

### Supplementary Fig. 4

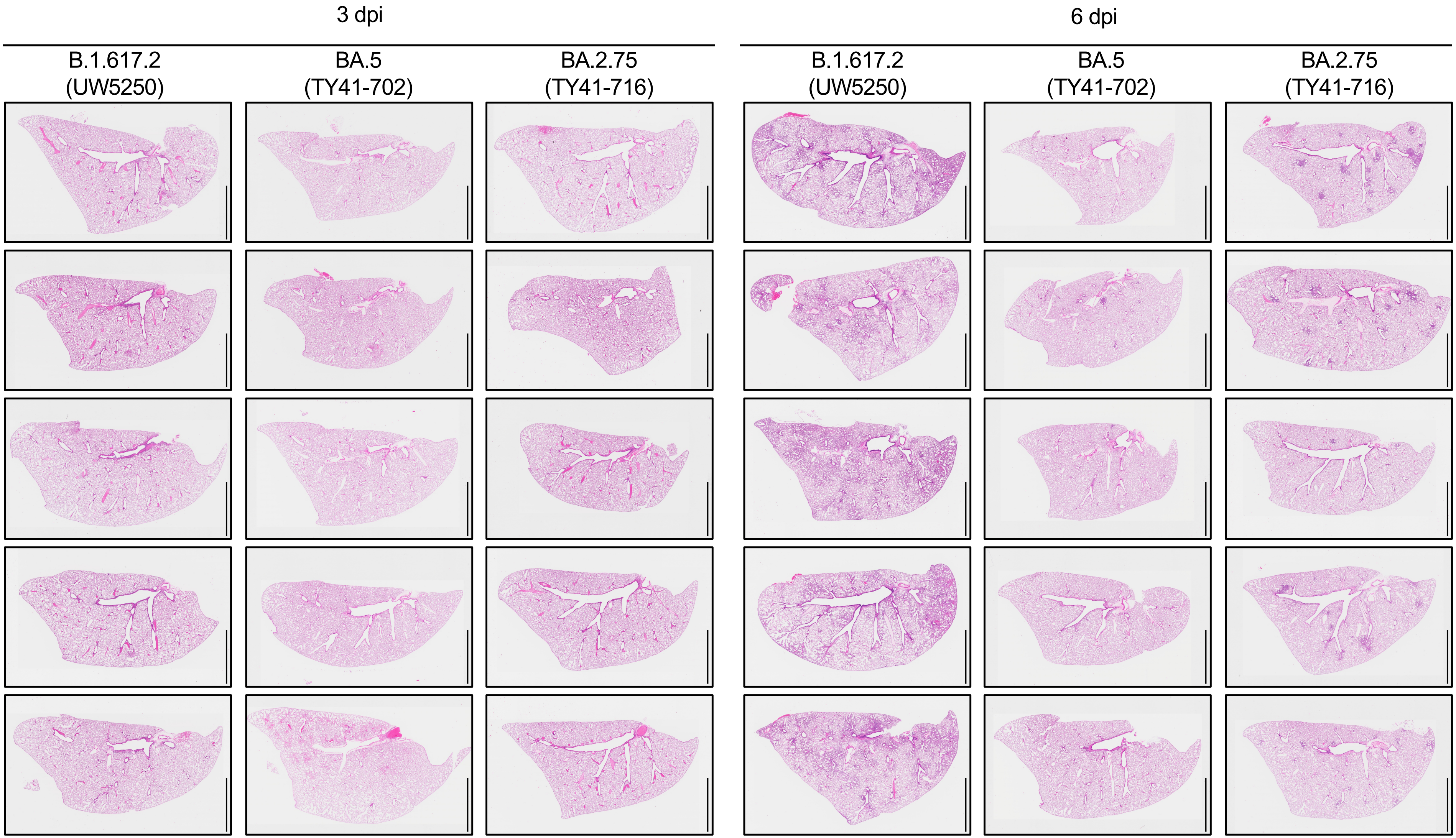

Supplementary Fig. 5

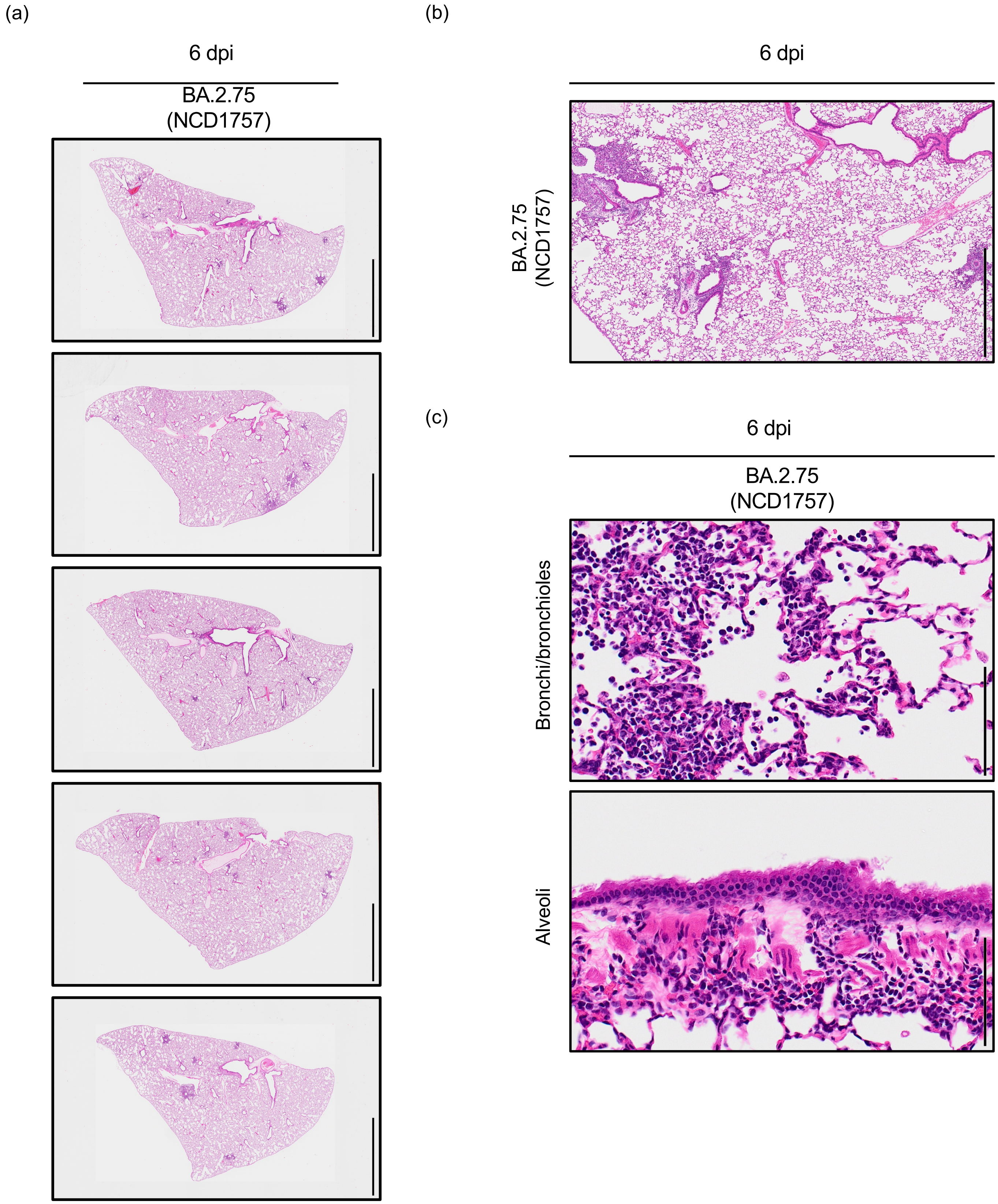
